# Attentional disengagement during external and internal distractions reduces neural speech tracking in background noise

**DOI:** 10.1101/2025.10.17.683146

**Authors:** Yue Ren, M. Eric Cui, Björn Herrmann

**Author notes:** Correspondence concerning this article should be addressed to Yue Ren and Björn Herrmann, Rotman Research Institute, Baycrest, 3560 Bathurst St, North York, ON, M6A 2E1, Canada. and.

## Abstract

Within-situation disengagement – the mental withdrawal during conversations in acoustically challenging environments – is a common experience of older people with hearing difficulties. Yet, most research on the neural mechanisms of attentional disengagement from speech listening has focused on the distraction by one competing speaker, whereas within-situation disengagement is often characterized by distraction towards external visual stimuli or internal thoughts and occurs in situations with ambient, multi-talker background masking. Across three electroencephalography (EEG) experiments in human participants of either sex, the current study examined how disengagement due to external and internal distractions affect the neural tracking of speech masked by different levels of multi-talker babble (speech in quiet, +6 dB, and −3 dB SNR). We observed enhanced early neural responses (<0.2 s) to the speech envelope for speech masked by background babble compared to speech in quiet (Experiments 1-3), suggesting stochastic facilitation. Importantly, neural tracking of the speech envelope was reduced when individuals were distracted by a visual-stimulus stream (Experiment 2) and by internal thought and imagination (Experiment 3). There were some indices suggesting the greatest disengagement-related decline in neural speech tracking occurs for the most difficult speech-masking condition, but this was not consistent across all measures. The current data show that disengagement due to external and internal distractions yield decreases in neural speech tracking, potentially suggesting converging neural pathways through which gain is downregulated in auditory cortex. These results indicate that disengagement from listening can be identified through non-invasive neural measures.

**Significance:** Many older adults with hearing difficulties mentally “tune out” during conversations in noisy environments, yet the neural mechanisms underlying this within-situation disengagement remain poorly understood. Across three electroencephalography experiments, we examined how external (visual) and internal (thought-based) distractions influence neural tracking of speech masked by multi-talker babble. We observed that attentional disengagement – whether induced by external stimuli or internal thoughts – reduced the brain’s tracking of the speech envelope. These findings demonstrate that listening disengagement can be objectively identified through neural measures and suggest a convergence in neural pathways through which both external and internal distractions down-regulate auditory gain, providing new insight into attentional control in challenging listening conditions.

## Introduction

Conversations in cafés or restaurants often involves listeners extracting meaning from a speaker amid background sounds. The cognitive resources required to comprehend under such conditions makes listening effortful (Pichora-Fuller et al., 2016; Peelle, 2018). When the listening situation becomes overly effortful, listeners may temporarily withdraw attention and “tune out” (Heffernan et al., 2016; McNeice et al., 2024; Sarant et al., 2024). This within-situation disengagement is a common response to sustained cognitive demand during listening (Heffernan et al., 2016; Herrmann and Johnsrude, 2020) and may promote social withdrawal (Motala et al., 2024). Characterizing markers of this disengagement could provide a window into how attention fluctuates during listening and towards detecting disengagement in real-time.

Individuals may disengage from speech listening for multiple reasons. Externally, attention may be captured by salient acoustic events, visual distractions, or competing speech inputs, disrupting the processing of a target stimulus (Lavie, 2005; Shinn-Cunningham, 2008). Internally, attention can drift away from the external environment as listeners engage in self-generated thought – often characterized as mind wandering – during which cognitive resources are reallocated toward memories, future plans, or imaginings (Eastwood et al., 2012; Ibaceta et al., 2024). This form of internally directed disengagement becomes more frequent when individuals listen to speech in background noise than when they listen in quiet (Steindorf et al., 2023). Thus, external distraction and internal drift represent two pathways to the same functional outcome of temporarily uncoupling attention from speech, but whether such disengagement leaves distinct neural signatures and whether such signatures depend on listening demands is unclear.

One increasingly common approach to investigate the encoding of continuous speech in the brain involves regressing relevant features of speech against signals recorded using electro- or magnetoencephalography (EEG; MEG; Lalor and Foxe, 2010; Ding and Simon, 2012; Hertrich et al., 2012). This approach, formalized as temporal response function (TRF) modeling (Crosse et al., 2016), is often used to quantify how neural activity tracks the temporal dynamics of the speech envelope, due to its importance for speech intelligibility (Ding et al., 2014; Lesenfants et al., 2019). The neural tracking of the speech envelope can be decreased when speech is masked by background sound (Ding and Simon, 2013; Synigal et al., 2023), although this seems to depend on the specific TRF metric and type of background masker, sometimes even leading to noise-related increases (Herrmann, 2025a, b).

Critically, a robust pattern across many studies is that the cortical tracking of the envelope of attended speech is stronger than the tracking of the envelope of speech that is ignored, reflecting selective amplification of behaviorally relevant neural signals (Kerlin et al., 2010; Presacco et al., 2016; Tune et al., 2021; Orf et al., 2025). Yet, most prior research on the effects of attention on speech tracking has relied on paradigms where a single talker competes with the target speech, whereas communication also often occurs in contexts with diffuse multi-talker background babble. Individuals may be particularly susceptible to attentional inward drifts under ambient background noise (Steindorf et al., 2023). Distraction-related disengagement in multi-talker settings need not resemble the explicit suppression of a competing speech stream used in selective-attention tasks. Because the target speech remains behaviorally relevant, a competing speaker is absent, and attentional decoupling may be intermittent, it is unclear whether external distraction or internal drift will produce measurable changes in speech tracking under multi-talker noise conditions.

We investigate how external and internal distractions affect neural speech tracking across varying levels of speech clarity using EEG and TRF analyses. After characterizing the dependence of neural speech tracking on speech clarity (Experiment 1), we examine how disengagement – from external visual distraction (Experiment 2) or internal thought (Experiment 3) – modulates neural tracking. We hypothesize that both forms of distraction will reduce neural speech encoding, particularly for noise-masked speech. Comparable effects across distraction types would point to a shared neurocognitive mechanism – such as down-regulation of auditory-cortical gain – through which attention decouples from sensory processing during effortful listening.

## General Methods

### Participants

Participants were aged 20-34 years, native English speakers or individuals who grew up in English-speaking countries (mostly Canada) and had been speaking English since early childhood (before age 6). All participants reported normal hearing abilities and no severe psychological or neurological disorders. Specific participant demographics are detailed for each experiment individually below.

Written informed consent was obtained prior to participation, and participants were compensated at a rate of $15 per hour. The study adhered to the Declaration of Helsinki, the Canadian Tri-Council Policy Statement on Ethical Conduct for Research Involving Humans (TCPS2-2014), and received ethical approval from the Research Ethics Board of the Rotman Research Institute at Baycrest Academy for Research and Education.

### Story materials and stimuli

In each of the three experiments, participants listened to ∼2-min stories. Stories were generated using OpenAI’s GPT-3.5 (OpenAI et al., 2023) based on researcher-provided prompts and manually reviewed to ensure unique character names, absence of non-English terms, proper use of punctuation, and avoidance of generic openings (e.g., “once upon a time”). Stories were revised manually or rewritten using GPT-3.5 as needed. Each story was on a different topic (e.g., high school reunion, friends on a trip; Panela et al., 2024; Herrmann, 2025a). For each story, GPT-3.5 was also used to generate five comprehension questions in a multiple-choice format, with four options per question. Comprehension questions and answers were manually edited wherever needed to ensure accuracy (Panela et al., 2024; Herrmann, 2025a).

Audio files were synthesized using Google’s AI-based speech synthesizer using the male voice en-US-Neural2-J (https://cloud.google.com/text-to-speech/docs/list-voices-and-types). Previous research has demonstrated that AI-generated speech is perceived as highly naturalistic and yields comparable speech-comprehension effects to human speech (Herrmann, 2023a, 2025c).

In each experiment, participants listened to 24 stories presented in 8 blocks (3 stories per block). Stories were presented under one of three listening conditions: clear speech (i.e., speech in quiet) or speech masked by 12-talker babble +6 dB SNR, or −3 dB SNR. Babble noise was derived from the masker materials of the Revised Speech in Noise test (Bilger, 1984). 12-talker babble simulates a crowded listening environment while preventing the identification of individual words in the masker (Bilger, 1984; Bilger et al., 1984; Wilson et al., 2012). The +6 dB and −3 dB SNR levels were chosen to create challenging listening conditions that correspond to approximately 90% and 55% speech intelligibility (Irsik et al., 2022). All stories and story-noise mixtures were normalized to the same root-mean-square (RMS) amplitude and presented at approximately 65 dB SPL.

### Acoustic presentation environment

Data collection took place in a sound-attenuating booth. Auditory stimuli were delivered via Sennheiser HD 25-SP II headphones, with audio output controlled by an RME Fireface UCX external sound card. Stimuli were presented using Psychtoolbox (v3.0.14) in MATLAB 2021b (MathWorks Inc.) on a Lenovo T480 laptop running Microsoft Windows 7. Visual stimuli were projected into the booth via screen mirroring.

### Electroencephalography (EEG) recording and preprocessing

EEG data were recorded from 32 scalp electrodes and the left and right mastoids using a BioSemi system (Amsterdam, The Netherlands). The online sampling rate was set to 1024 Hz, with a low-pass filter of 208 Hz. Electrodes were referenced online using a monopolar reference feedback loop, which connected a driven passive sensor and a common-mode-sense (CMS) active sensor, both positioned posteriorly on the scalp.

Preprocessing was conducted in MATLAB (MathWorks, Inc.). Data were first notch-filtered to suppress 60-Hz power line noise using an elliptic filter. Signals were then re-referenced to the averaged mastoids to enhance the signal-to-noise ratio for auditory responses at fronto-central electrodes (Ruhnau et al., 2012; Herrmann, 2023b). Additional filtering included a 0.7 Hz high-pass filter (2449 samples, Hann window) and a 22 Hz low-pass filter (211 samples, Kaiser window with β = 4). EEG data were segmented into time series locked to story onset and downsampled to 512 Hz. Artifact removal was performed using Independent Component Analysis (ICA) in FieldTrip MATLAB toolbox to eliminate blinks, eye movements, and muscle artifacts (Makeig et al., 1995; Oostenveld et al., 2011). After ICA, further artifact rejection was applied by zeroing segments where EEG amplitude varied by more than 80 μV within 0.2 s in any channel (Dmochowski et al., 2012; Cohen and Parra, 2016). Finally, data were low-pass filtered at 10 Hz (251 points, Kaiser window) to focus on low-frequency neural signals most sensitive to acoustic features (Di Liberto et al., 2015; Zuk et al., 2021).

### Calculation of amplitude-onset envelopes

For each clear speech signal, a cochleogram was generated using a simple auditory-periphery model consisting of 30 auditory filters (McDermott and Simoncelli, 2011). The amplitude envelope obtained for each auditory filter was compressed by a factor of 0.6 to simulate inner ear compression (McDermott and Simoncelli, 2011). The amplitude envelopes were then averaged across auditory filters and low-pass filtered using a 40-Hz Butterworth filter. The first derivative was calculated and all negative values were set to zero to obtain the amplitude-onset envelope (Hertrich et al., 2012; Tune et al., 2021; Yasmin et al., 2023; Panela et al., 2024). The amplitude-onset envelope was computed since it elicits strong neural speech tracking (Hertrich et al., 2012). The amplitude-onset envelope was down-sampled to match the EEG sampling rate.

### Analysis of neural tracking of speech

To investigate neural speech tracking, we adopted Temporal Response Functions (TRFs) and assessed EEG prediction accuracy (Crosse et al., 2016; Crosse et al., 2021). The TRF is a system identification model that employs regularized linear regression to relate sensory stimulus features with neural responses. Specifically, we utilized the Multivariate Temporal Response Function (mTRF) Toolbox in MATLAB (Crosse et al., 2016; Crosse et al., 2021) to map the amplitude-onset envelope of speech onto EEG signals (forward model). To mitigate overfitting, ridge regularization was applied, with the ridge parameter (λ) set to 10, a value informed by prior research to avoid extreme value selection during cross-validation and optimize computational efficiency (Fiedler et al., 2017; Fiedler et al., 2019; Yasmin et al., 2023; Panela et al., 2024).

For each story, 100 randomly selected 25-s EEG snippets and their corresponding time series of the amplitude-onset envelope were extracted for model training and testing using leave-one-sample-out cross-validation. On each iteration, one snippet’s EEG data and the corresponding time series of the amplitude-onset envelope were reserved as the test set, while the remaining, non-overlapping data were used for training. The model was trained using time lags ranging from 0 to 0.4 s, employing the *mTRFcrossval* and *mTRFtrain* functions. To assess prediction accuracy, the acoustic feature time series of the test dataset was convolved with the TRF weights to generate a predicted EEG time series, separately for each of the 100 iterations. EEG prediction accuracy was calculated as the Pearson correlation between the predicted and actual EEG data, computing using the *mTRFpredict* function. EEG prediction accuracy was averaged across the leave-one-out iterations.

To investigate the neural-tracking response amplitude at a more nuanced timescale, TRFs for each training dataset were calculated for lags ranging from −0.15–0.55 s. Baseline correction was performed by subtracting the mean signal across −0.15–0 s from the TRF data at each time point. The weights were averaged across the 100 training datasets.

For each participant, TRF weights and EEG prediction accuracy were averaged across stories, separately for the three different speech-clarity conditions and averaged across the frontal-central electrode cluster (F3, F4, Fz, FC1, FC2, Cz, FC5, FC6), which is sensitive to auditory cortical activity (Näätänen and Picton, 1987; Picton et al., 2003).

Time courses of TRF weights exhibit positive and negative deflections (peaks), akin to traditional event-related potentials (Luck, 2014; Crosse et al., 2016; Crosse et al., 2021). To examine response magnitude differences across speech-clarity levels and attention conditions, we focused on the P1-N1 (first major positive peak minus first major negative peak) and P2-N1 amplitude differences. Using peak-to-peak measures helps reduce sensitivity to slow baseline shifts and condition-related waveform morphology differences, providing a robust index of response magnitude across conditions. The amplitudes of P1, N1, and P2 components were determined by local extrema (maxima for P1 and P2, minima for N1) within a 0.1-s time window centered on each component based on averaged data across all participants in each experiment. Latencies for P1, N1, and P2 were identified as the time points corresponding to these extrema. For each participant, amplitudes were calculated within a 0.03-s window (±0.015 s) centered on the latencies of each component from grand-average data. This approach accommodates inter-individual latency jitter and avoids unreliable participant-specific peak-picking in conditions where some participants might not show a clear extremum, while keeping the P1, N1, and P2 windows non-overlapping. The P1-N1 and P2-N1 amplitude differences were computed by subtracting N1 amplitudes from P1 and P2, respectively.

We also explored the extent to which semantic surprisal was tracked by the brain, but these analyses were ambiguous and did not enable drawing clear conclusions about the semantic processing in the current study, mirroring to some extent other debates in the field (Daube et al., 2019; Gillis et al., 2021; Verschueren et al., 2022; Chalehchaleh et al., 2025). We provide these explorations in the Supplementary Materials (Table S1, Figures S1 and S2).

### Statistical analyses

All statistical analyses were carried out using MATLAB (MathWorks) and JASP software (version 0.19.2). Details about the specific tests used are provided in the relevant sections for each experiment.

## Experiment 1: Influences of speech clarity on neural speech tracking

Experiment 1 measured speech-evoked EEG activity under focused listening across speech-clarity conditions to establish the baseline signal changes underlying distraction effects tested in Experiments 2 and 3.

### Method and materials

#### Participants

Twenty-four younger adults participated in Experiment 1 (mean = 25.2 years; SD = 3.85; 14 females, 10 males).

#### Stimuli and procedure

Twenty-four stories, each lasting between 109 and 133 seconds, were generated for Experiment 1. The stories were presented under one of three speech-clarity conditions: clear speech (no noise), +6 dB SNR, or −3 dB SNR (12-talker babble). The experiment was divided into 8 blocks, with each block containing 3 stories with the 3 speech-clarity conditions. Assignment of speech-clarity conditions to specific stories was counter-balanced across participants. During each trial, participants listened attentively to the story while their EEG signals were recorded. After each story, participants answered five multiple-choice comprehension questions, each with four response options (chance level = 25%). Additionally, participants provided a self-reported gist rating on a scale from 1 to 9, with higher ratings indicating a better understanding of the story’s gist. Gist ratings correlate highly with speech intelligibility as measured using word report scores (Davis and Johnsrude, 2003; Ritz et al., 2022). The exact wording for the gist rating was: “I understood the gist of the story. Please rate this statement independently of how you felt about the other stories”. Gist ratings were linearly transformed to ranging from 0 to 1 in order to facilitate interpretation similar to proportions (Mathiesen et al., 2024; Herrmann, 2025a). Participants underwent a practice block prior to the main experimental procedures. The full experimental session lasted approximately 2 hours per participant, with the experimental procedures – including EEG recording – taking about 1 hour.

#### Analysis

The effect of Speech Clarity (clear, +6 dB, −3 dB SNR) on comprehension accuracy, gist rating, EEG prediction accuracy, P1-N1 amplitude, and P2-N1 amplitude were evaluated in separate one-way repeated-measures analysis of variance (rmANOVA). Post-hoc pairwise comparisons were performed using the Holm correction for multiple comparisons (Holm, 1979).

## Results

Both comprehension accuracy and gist ratings (Figure 1A) were affected by speech clarity (comprehension: F_2, 46_ = 11.986, p = 6.5·10^-5^, η_p_^2^ = .343; gist: F_2, 46_ = 34.466, p = 7.1·10^-10^, η_p_^2^ = .600). Performance dropped for the lowest speech clarity (−3 dB SNR): pair-wise comparisons showed lower scores at −3 dB than at clear and at +6 dB for both behavioural measures (all p_Holm_ < .01). Clear and +6 dB did not differ significantly on either measure (all p_Holm_ > .05).

**Figure 1:**
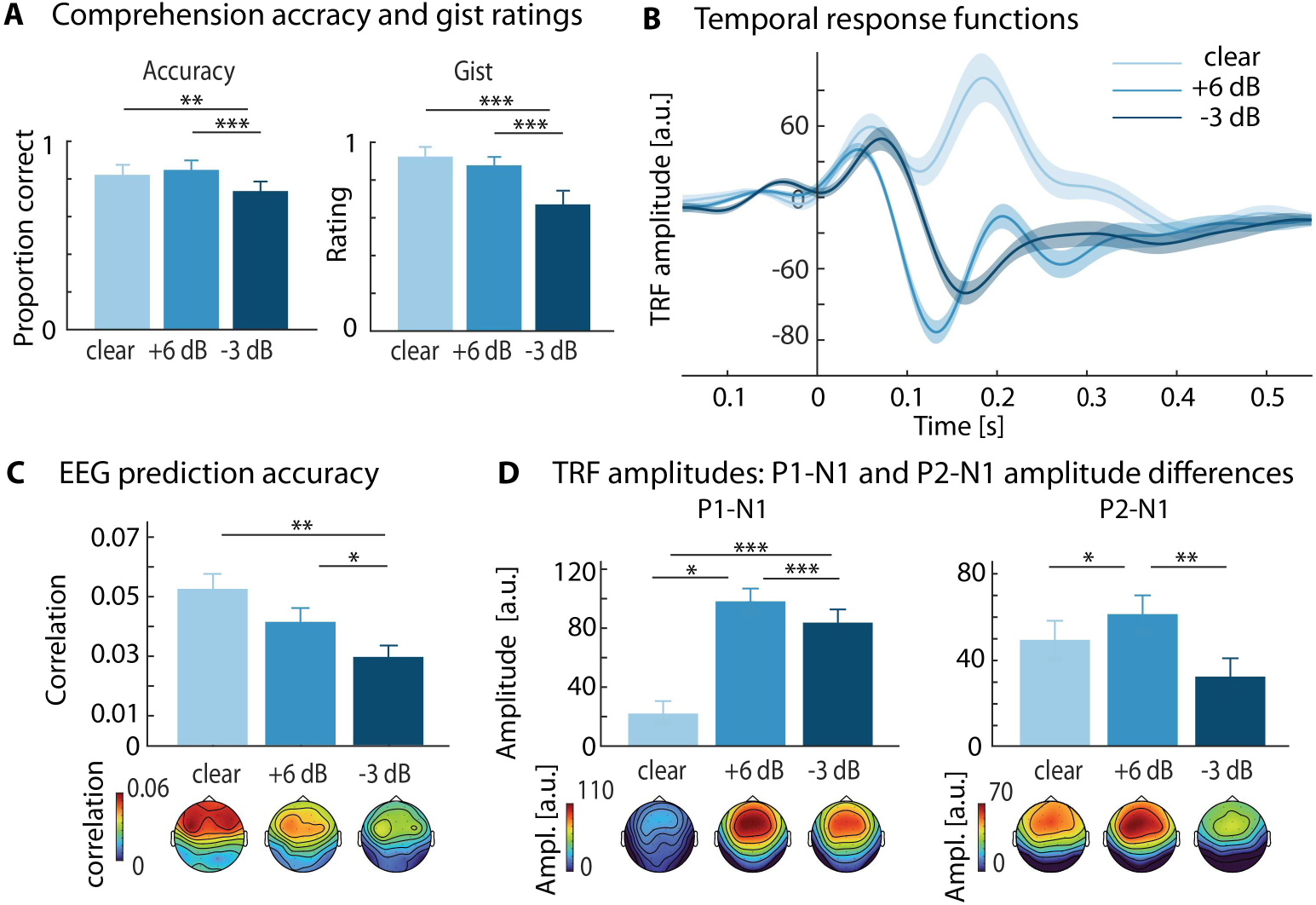
Results for Experiment 1. **A:** Comprehension scores (left), gist ratings (right) across different speech-clarity conditions. **B:** Temporal response functions (TRF) waveforms. **C:** EEG prediction accuracy and corresponding topographies. **D:** P1-N1 (left) and P2-N1 (right) amplitude and topographies for each speech-clarity condition. Error bars denote the standard error of the mean. *p < .05, **p < .01, ***p < .001 (Holm-corrected where applicable).

EEG prediction accuracy (Figure 1C) showed a significant main effect of speech clarity (F_2, 46_ = 9.376, p = 3.8·10^-4^, η_p_^2^ = .269). The values were lower at −3 dB SNR compared to clear (t_23_ = 3.882, p_Holm_ = .002, d = .843) and +6 dB (t_23_ = 2.843, p_Holm_ = .018, d = .471), whereas clear and +6 dB did not differ (t_23_ = 1.966, p_Holm_ = .062, d = .409). Scalp topographies revealed highest EEG prediction accuracy at frontal-central electrodes that are sensitive to auditory cortex activity.

TRF time courses are displayed in Figure 1B, indicating that background noise affects the shape and deflections of the response. To avoid that analyses of response amplitudes are affected by overall morphological differences, analyses focused on the P1-N1 and P2-N1 amplitudes (Lanting et al., 2013; de Boer and Krumbholz, 2018; Herrmann, 2025a, b). The speech-clarity effect for the P1-N1 amplitude (F_2, 46_ = 52.108, p = 1.5·10^-12^, η_p_^2^ = .694) revealed a non-monotonic pattern: responses were largest at +6 dB and remained elevated at −3 dB relative to clear (all p_Holm_ < .05). For the P2-N1 amplitude the effect of speech clarity (F_2, 46_ = 8.110, p = 9.6·10^-4^, η_p_^2^ = .261) showed larger amplitudes at +6 dB compared to −3 dB and clear (both p_Holm_ < .05), whereas −3 dB and clear did not differ significantly (p_Holm_ = .110; Figures 1B, D).

In sum, the results of Experiment 1 show that background babble impairs speech comprehension, gist rating, and overall EEG prediction accuracy, with declines observed only at the lowest speech clarity (−3 dB). Early TRF responses (both P1-N1 and P2-N1 components) were selectively enhanced under moderate noise – peaking at + 6 dB – consistent with recent work suggesting stochastic resonance phenomena in neural speech tracking and for more simple sounds (Parbery-Clark et al., 2011; Alain et al., 2014; Herrmann, 2025a, b). With the foundation of Experiment 1, Experiment 2 examines how disengagement from listening due to visual distraction affects neural speech tracking, and whether disengagement effects are affected by speech clarity.

## Experiment 2: Influences of visual distraction on neural speech tracking

### Method and materials

#### Participants

Twenty-four younger adults participated in Experiment 2 (mean = 26 years; SD = 3.95; 14 females, 10 males).

#### Stimuli and procedure

We generated a new set of 24 stories (102–141 s each) for Experiment 2. Similar to Experiment 1, the session comprised 8 blocks, with each block containing 3 stories, one for each speech-clarity condition (clear, +6 dB, and −3 dB SNR). In four blocks, participants attended to the speech (‘attend’ condition), whereas in the other four blocks, participants ignored the speech (‘ignore’ condition). Assignment of speech clarity and attention conditions to specific stories was counter-balanced across participants. Blocks alternated in attention conditions, and the starting attention condition was counter-balanced across participants.

In ‘Ignore’ blocks, participants were asked to ignore the stories and instead perform a visual 1-back task. The visual 1-back task utilized images of white digits (0 to 9) on a black background, sourced from the MNIST Handwritten Digit Classification Dataset (Herrmann 2024; Deng et al. 2012). Each digit image was displayed for 0.25 seconds, followed by a 0.25-s black screen before the next image appeared. Visual stimuli thus repeat as a rate of 2 Hz. The digit stream included a digit repetition – albeit a different image sample – every 6 to 12 digits. Participants were asked to press a keyboard button as soon as they detected a repetition. Hit rates and response times were recorded. Additionally, participants rated the degree to which they were able to ignore the speech using a 9-point scale (1 = strongly disagree, 9 = strongly agree; linearly transformed to range from 0-1). No comprehension questions or gist ratings were used in the ‘Ignore’ blocks to avoid incentivizing participants to listen to the stories. In ‘Attend’ blocks, participants were instructed to listen attentively to the stories, followed by comprehension questions and a gist rating (similar to Experiment 1). Visual digit images at a 2-Hz rate were presented concurrently with the stories to ensure similar visual and auditory stimulus presentation in ‘attend’ and ‘ignore’ blocks, but participants were asked to pay attention only to the speech in ‘attend’ blocks and ignore the visual stimuli.

Participants underwent a practice block prior to the main experimental procedures to familiarize them with the tasks. The full experimental session lasted approximately 2 hours per participant, with the experimental procedures – including EEG recording – taking about 1 hour.

#### Analysis

For each participant, comprehension accuracy and gist ratings from ‘attend’ blocks were averaged across the four stories per speech-clarity condition (clear, +6 dB, −3 dB). Reaction times, visual detection accuracy, and able-to-ignore ratings from ‘ignore’ blocks were also averaged across the four stories per speech-clarity condition. For each behavioral measure, we calculated a one-way rmANOVA with the within-participant factor Speech Clarity (clear, +6 dB, −3 dB). Note that behavioral measures did not comprise the Attention factor.

Because Experiment 2 comprised a repetitive visual stream of digits occurring at 2 Hz, this provided an opportunity to neurally investigate attention to the visual stream. To this end, EEG data for each story / visual steam were divided into 7-s epochs (shifted by 0.5 s across the story duration) that were time-locked to a visual stimulus. A fast Fourier transform (FFT) was calculated for each epoch. Inter-trial phase coherence (ITPC; Lachaux et al., 1999) was then calculated across epochs to obtain a measure of neural responses to the visual stream. ITPC was used here instead of evoked amplitude because ITPC spectra do not show 1/f patterns that amplitude spectra exhibit (Herrmann et al., 2019; Keitel et al., 2019; Irsik et al., 2021), but which can complicate analyses (Donoghue et al., 2020; Donoghue et al., 2022). Mathematically, ITPC and evoked amplitude are otherwise closely related. We calculated the average across ITPC at the 2-Hz visual repetition rate and its harmonics up to 10 Hz (Irsik et al., 2021). Taking into account the harmonics is needed because data are not sinusoidal in the time domain (as indicated by the presence of harmonics). ITPC was averaged across 3 occipital channels (O1, Oz, O2) to focus on visual-cortex activity. For the statistical analysis, ITPC was averaged across stories, separately for each speech-clarity condition (clear, +6 dB, −3 dB) and attention condition (Attend, Ignore), and a rmANOVA with the two factors was calculated.

For the neural speech tracking analysis, EEG prediction accuracy, P1-N1 amplitude, and P2-N1 amplitude were evaluated in separate rmANOVAs with the factors Speech Clarity (clear, +6 dB, −3 dB) and Attention (Attend, Ignore). Post-hoc pairwise comparisons were performed using the Holm correction for multiple comparisons (Holm, 1979).

## Results

Behavioral performance during ‘attend’ speech blocks are shown in Figure 2A. Speech Clarity affected both comprehension accuracy (F_2, 46_ = 11.279, p = 1·10^-4^, η_p2_ = .329) and gist ratings (F_2, 46_ = 47.964, p = 5.6·10^-12^, η_p2_ = .676). Similar to Experiment 1, performance was lower in the −3 dB condition compared to the clear and +6 dB conditions for both measures (both p_Holm_ < .01). For gist rating, +6 dB was also lower than clear (t_23_ = 3.174, p_Holm_ = .004, d = .444), whereas for comprehension accuracy, clear and +6 dB did not differ (t_23_ = .554, p_Holm_ = .585, d = .103). During ‘ignore’ blocks (Figure 2B), speech clarity did not affect visual-detection accuracy (F_2, 46_ = .506, p = .606, η_p2_ = .022) or reaction times (F_2, 46_ = 2.855, p = .068, η_p2_ = .110). In contrast, ‘able-to-ignore’ ratings indicate that participants found the stories at −3 dB easier to ignore compared to clear and +6 dB (both p_Holm_ < .05; effect of Speech Clarity: F_2, 46_ = 7.828, p = .001, η_p_^2^ = .254).

**Figure 2:**
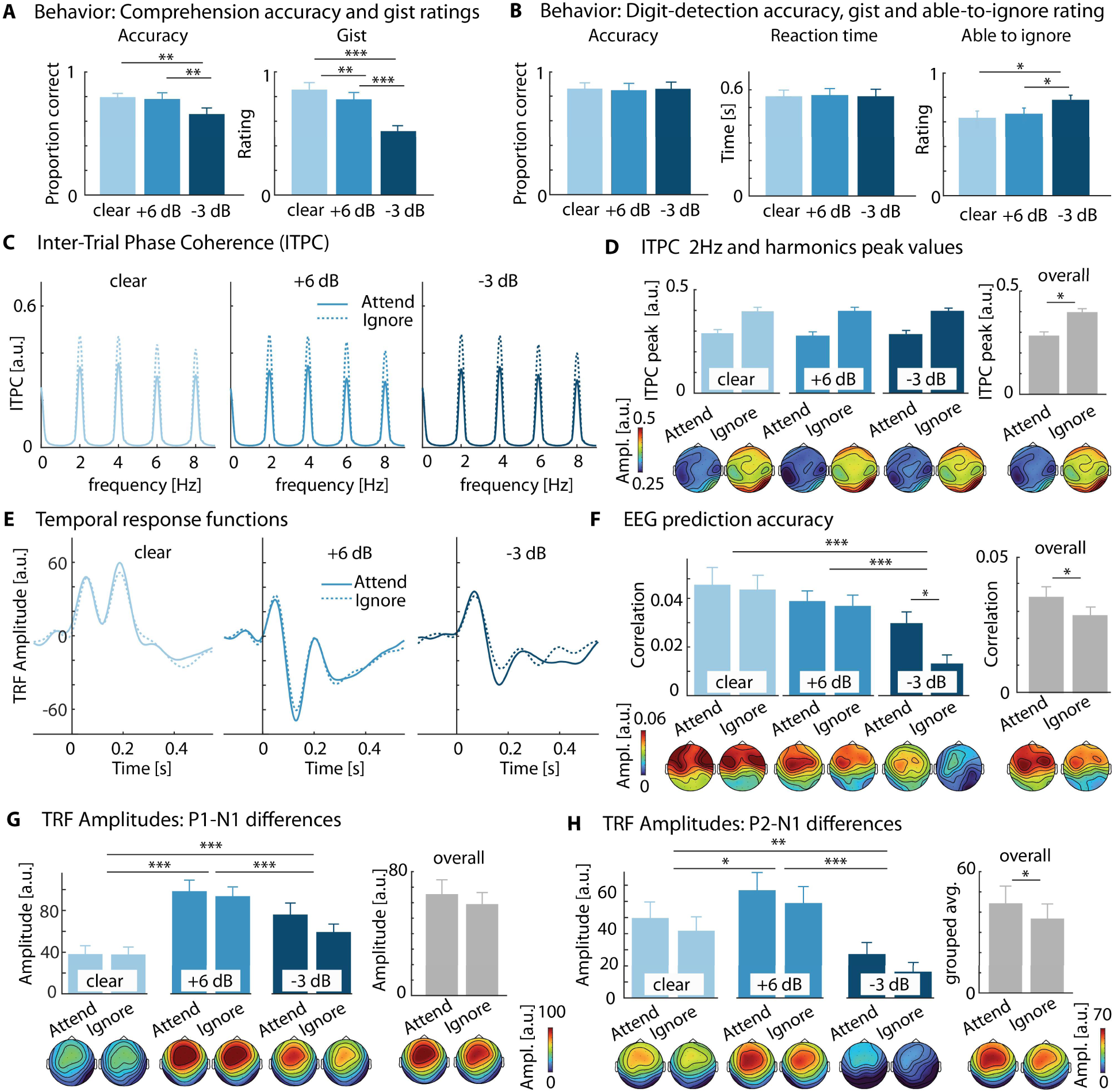
Results for Experiment 2. **A:** Comprehension scores (left) and gist ratings (right) across three speech-clarity levels during ‘attend’ blocks. **B:** Detection accuracy and reaction times (RT) for the visual 1-back task, and ratings of the ability to ignore the spoken stories during visual distraction blocks. **C:** ITPC showing peaks at 2 Hz and harmonics, reflecting responses to the visual stimuli occurring at a 2-Hz stimulation rate. **D:** Peak ITPC (2-10 Hz averaged) for three speech-clarity conditions separated by attention blocks. **E:** Temporal response functions (TRFs) for ‘attend’ (solid) and ‘ignore’ (dashed) conditions for the different speech-clarity conditions. **F:** Mean EEG prediction accuracy and corresponding topographies. **G:** Mean P1-N1 amplitudes and corresponding topographies. **H:** Mean P2-N1 amplitudes and corresponding topographies. Error bars denote the standard error of the mean. *p < .05, **p < .01, ***p < .001 (Holm-corrected where applicable).

Consistent with the periodic visual stream and isochronously occurring visual evoked potentials, ITPC showed clear peaks at 2 Hz and harmonics (Figure 2C). Peak ITPC (2–10 Hz harmonics averaged; Figure 2D) was higher when participants ignored than attended to the speech (F_1, 23_ = 78.844, p = 6.8·10^-9^, η_p_^2^ = .774). There was no effect of Speech Clarity (F = .882, p = .421, η_p_^2^ = .037), and no interaction (F_2, 46_ = 1.676, p = .198, η_p_^2^ = .068). These data show that participants indeed attended to the visual digit stream more when they ignored relative when they attended to the speech.

EEG prediction accuracy was lower for −3 dB than at Clear (t_23_ = 4.137, p_Holm_ = 8·10^-4^, d = .882) and +6 dB (t_23_ = 4.665, p_Holm_ = 3.2·10^-4^, d = .689), with no difference between Clear and +6 dB (t_23_ = 1.001, p_Holm_ = .327, d = .193; effect of Speech Clarity: F_2, 46_ = 12.368, p = 5·10^-5^, η_p_^2^ = .350 Figure 2F). EEG prediction accuracy was greater for the attend than ignore conditions (effect of Attention: F_1, 23_ = 7.244, p = .013, η_p_^2^ = .240), although the significant interaction (F_2, 46_ = 3.561, p = .036, η_p_^2^ = .134) shows that this difference was specific for the −3 dB SNR condition (t_23_ = 3.346, p_Holm_ = .031, d = .703).

TRF time courses are displayed in Figure 2E. The P1-N1 amplitude was smallest for clear speech, largest for +6 dB SNR, and in-between for −3 dB (all pairwise p_Holm_ < .001; effect of Speech Clarity: F_2, 46_ = 33.186, p = 1.2·10^-9^, η_p_^2^ = .591). There was no effect of Attention (F_2, 46_ = 3.008, p = .096, η_p_^2^ = .116), but the interaction between Speech Clarity and Attention was significant (F = 3.726, p = .032, η_p_^2^ = .139). Although not surviving Holm correction, the interaction was driven by the attention effect at −3 dB SNR as indicated by the uncorrected comparison (t_23_ = 2.471, p = .021, p_Holm_ = .102, d = .504). Attention effects for other the two speech-clarity levels were not significant (both effects, p_Holm_ and p > .50).

The P2-N1 amplitude was larger at +6 dB than at clear (t_23_ = 2.648, p_Holm_ = .014, d = .382) and −3 dB (t_23_ = 6.342, p_Holm_ = 5.4·10^-6^, d = .947), and amplitudes for clear speech were larger than for −3 dB (t_23_ = 3.450, p_Holm_ = .004, d = .565; effect of Speech Clarity: F_2, 46_ = 19.481, p = 7.4·10^-7^, η_p2_ = .459). P2-N1 amplitudes were larger when participants attended to the speech compared to when they ignored it (effect of Attention: F_2, 46_ = 4.426, p = .047, d = .194). There was no interaction (F_2, 46_ = 0.089, p = .915, η_p2_ = .004).

In sum, disengagement from speech listening due to visual distraction reduced signatures of neural speech tracking (EEG prediction accuracy and TRF component amplitudes), most prominently for speech masked at −3 dB SNR. This result was mirrored by participants also reporting greater ease ignoring speech at −3 dB than at +6 dB and clear speech. Experiment 2 demonstrates that neural speech tracking is sensitive to disengagement from listening due to external distraction, such that the amplitude-envelope onset of the speech is encoded less strongly. Critically, disengagement from listening in everyday life not only occurs because of external distraction, but also because of internal distraction when a listener’s mind is wandering off. Experiment 3 was designed to examine whether disengagement due to internal distractions affects neural speech tracking.

## Experiment 3: Influences of internal imagination on neural speech tracking

### Method and materials

#### Participants

Twenty-four younger adults were recruited for Experiment 3 (mean = 26.3 years, SD = 4.37; 13 females, 10 males, 1 non-binary).

### Stimuli and procedure

A new set of 24 stories was generated, each lasting between 111 and 139 seconds. The experiment was divided into 8 blocks, with each block containing 3 stories, one for each speech-clarity condition (clear, +6 dB, and −3 dB SNR). In four blocks, participants attended to the speech (‘attend’ condition), whereas in the other four blocks, participants ignored the speech (‘ignore’ condition). Assignment of speech clarity and attention conditions to specific stories was counter-balanced across participants. Blocks with different attention conditions alternated in the experimental session and starting attention condition in the session was counter-balanced across participants. Participants underwent a practice block prior to the main experimental procedures to familiarize them with the tasks.

In the ‘attend’ blocks, participants attentively listen to the stories, answered comprehension questions, and rating gist understanding (similar to Experiments 1 and 2). In ‘ignore’ blocks, participants were asked to disregard the spoken stories and engage in an ‘imagination’ task, where they mentally wandered off. During the imagination task, participants were provided with a verbal instruction outlining a daily scenario accompanied by three concrete cues (e.g., scenario: public transport commute; cues: seats, tapping cards, and bus driver). After a brief preparation period, participants were instructed to immerse themselves in the imagined scenario as soon as the auditory stimulus began. They were encouraged to use internal narratives and mental imagery to facilitate the task, and to mentally jump freely to any other scenarios that come to mind as much as they wanted. This task aimed to mirror disengagement in daily life where people may think about concrete daily activities or interactions (Smallwood and Andrews-Hanna, 2013; Ibaceta et al., 2024), although here facilitated by task instructions. Following each story, participants evaluated their experience using a 9-point scale (1 = strongly disagree, 9 = strongly agree) across four statements: whether they were able to ignore the speech, mentally wander off, visualize the given scenario, and would be able retell the story. No comprehension questions or gist ratings were used in the ‘ignore’ blocks to avoid that participants attend to the stories. Ratings were linearly transformed to range from 0-1 to facilitate interpretability (Mathiesen et al., 2024; Herrmann, 2025a).

Participants underwent a practice block prior to the main experimental procedures to familiarize them with the tasks. The full experimental session lasted approximately 2 hours per participant, with the experimental procedures – including EEG recording – taking about 1 hour.

#### Analysis

For each behavioral measure, a one-way rmANOVA with the within-participant factor Speech Clarity was calculated. Behavioral measures did not comprise the Attention factor to avoid that participants are biased towards the irrelevant dimension. For each EEG measure (EEG prediction accuracy, P1-N1 amplitude, and P2-N1 amplitude), a two-way rmANOVA was conducted using the with-participant factors Speech Clarity (clear, +6 dB, and −3 dB SNR) and Attention (Attend, Ignore).

## Results

Speech clarity affected both comprehension accuracy (F_2, 46_ = 20.647, p = 4·10^-7^, η_p_^2^ = .473) and gist ratings (F_2, 46_ = 40.847, p = 7.5·10^-11^, η_p_^2^ = .637). Both behavioral measures were lower at −3 dB compared to the clear (both p_Holm_ < .001) and +6 dB (both p_Holm_ < .001), with no difference between Clear and +6 dB (both p_Holm_ > .05; Figure 3A).

**Figure 3:**
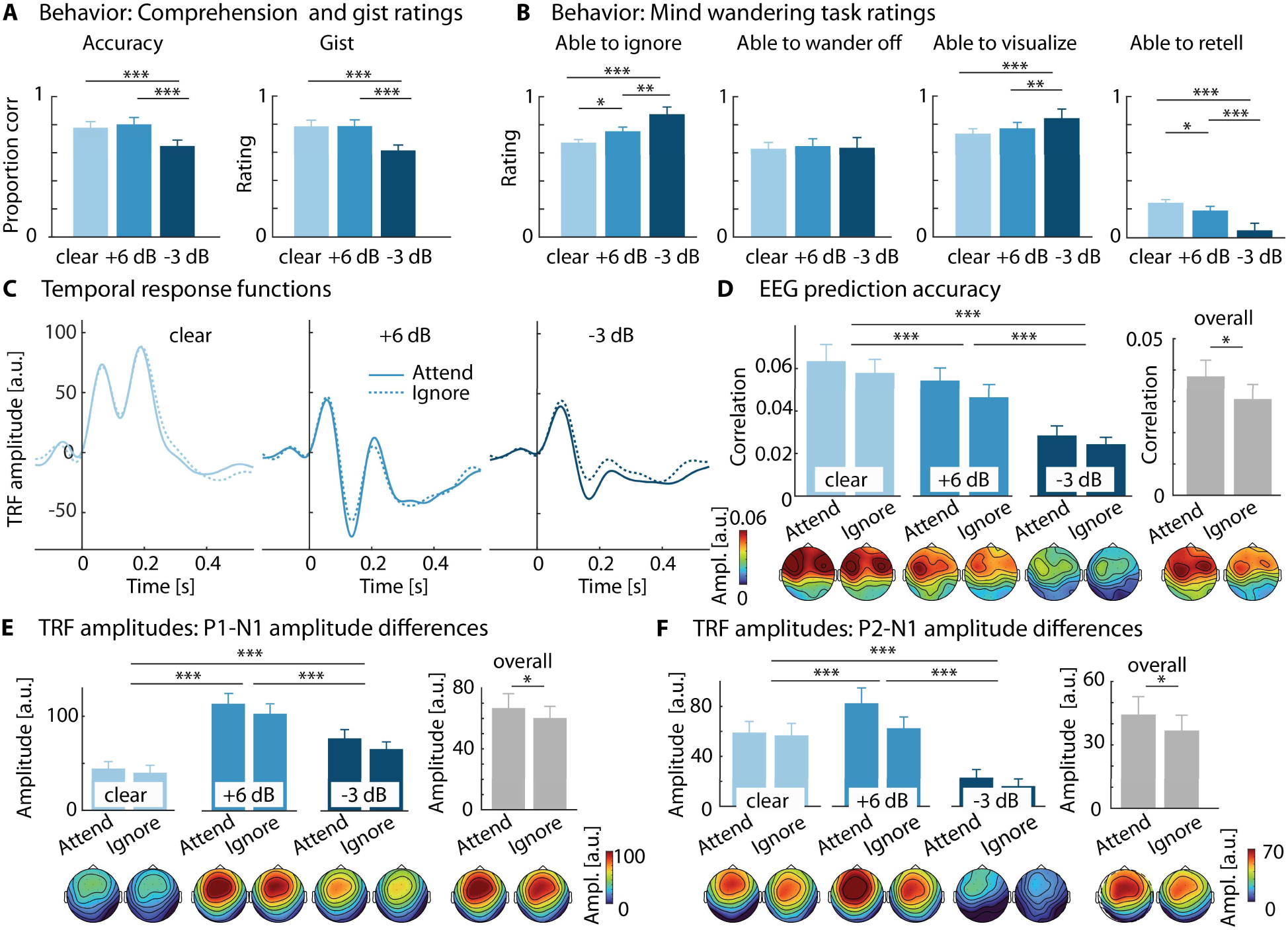
Results for Experiment 3. **A:** Comprehension scores (left) and gist ratings (right) across three speech-clarity conditions during attentive listening blocks. **B:** Self-evaluations of the ability ‘to ignore the spoken stories’, ‘to be able to wander off’, ‘to be able to visualize the given scenery’, ‘to be able to retell the spoken stories’ during imagination blocks. **C:** Temporal response functions (TRFs) during Attend (solid) and Ignore (dashed) across Speech Clarity. **D:** EEG prediction accuracy and corresponding topographies. **E:** P1-N1 amplitudes and corresponding topographies. **F:** P2-N1 amplitudes and corresponding topographies. Error bars denote the standard error of the mean. *p < .05, **p < .01, ***p < .001 (Holm-corrected where applicable).

Behavioral ratings related to the imagination task are shown in Figure 3B. Participants rated it to be easier to ignore stories (effect of Speech Clarity: F_2, 46_ = 18.591, p = 1.2·10^-6^, η_p2_ = .447; p_Holm_ < .05 for all pairwise comparisons) and to be less able to retell stories with decreasing speech clarity (effect of Speech Clarity: F_2, 46_ = 26.878, p = 1.9·10^-8^, η_p2_ = .539; p_Holm_ < .05 for all comparisons). The ability to mentally visualize was rated higher at −3 dB compared to Clear (t_23_ = 4.236, p_Holm_ = 9.4·10^-4^, d = .552) and +6 dB (t_23_ = 3.636, p_Holm_ = .003, d = .364; effect of Speech Clarity: F_2, 46_ = 11.030, p = 1.2·10^-4^, η_p_^2^ = .324), but speech clarity had no effect on participants’ ability to wander off mentally (F_2, 46_ = .215, p = .807, η_p_^2^ = .009). Overall, the ratings suggest that participants could well imagine mentally and ignore the speech, such that they felt relatively unable to retell the story.

EEG prediction accuracy decreased with decreasing speech clarity (F_2, 46_ = 26.774, p = 1.9·10^-8^, η_p_^2^ = .538; p < .05 for all pairwise comparisons; Figure 3D). Critically, EEG prediction accuracy was lower during imagination (ignore) than attentive listening (effect of Attention: F_1, 23_ = 7.405, p = .012, η_p_^2^ = .244), with no Speech Clarity × Attention interaction (F =.091, p = .913, η_p_^2^ = .004).

TRF time courses are depicted in Figure 3C. Speech Clarity modulated the P1-N1 amplitudes (main effect of Speech Clarity: F_2, 46_ = 40.196, p 8·10^-11^, η_p_^2^ = .636), as amplitudes were largest at +6 dB and smallest in the Clear condition (all pairwise p_Holm_ < .001). Mental imagination reduced P1-N1 amplitudes compared to attentive speech listening (effect of Attention: F_1, 23_ = 4.862, p = .038, η_p_^2^ = .175). The interaction was not significant (F = .585, p = .561, η_p_^2^ = .025).

P2-N1 amplitudes also varied with Speech Clarity (F_2, 46_ = 29.536, p = 5.6·10^-9^, η_p_^2^ = .562), such that amplitudes were largest at +6 dB and smallest in the Clear condition (all pairwise p_Holm_ < .05). Mental imagination reduced P2-N1 amplitude compared to attentive speech listening (effect of Attention: F_1, 23_ = 5.837, p = .024, η_p_^2^ = .202). The interaction was not significant (F_2, 46_ = 1.516, p = .230, η_p_^2^ = .062).

In sum, Experiment 3 shows, consistent with the findings from Experiments 1 and 2, that background babble can enhance early neural responses to the amplitude-envelope of speech (see also Yasmin et al., 2023; Panela et al., 2024; Herrmann, 2025a). Experiment 3 results further indicate that participants can ignore speech well while engaging in a mental imagination task. Critically, Experiment 3 extends the external distraction effect in Experiment 2 by showing that internal distraction (imagination) reduces the encoding of the amplitude-envelope of speech, indicated by all three signatures of neural speech tracking. The main effect of attention in the absence of an interaction indicates that internal distraction reduces speech encoding across speech-clarity levels.

### Comparison of attention effects between external and internal distraction

In order to directly compare the attention effects between external and internal distraction, we first calculated an rmANOVA for the participants’ ratings of their ability to ignore speech when they performed the visual distraction or internal imagination task. Rating scores were used as dependent measure, with the within-participant factor Speech Clarity (clear, +6 dB, −3 dB) and the between-participants factor Distraction Type (external, internal; Experiments 2 and 3). The rmANOVA showed higher speech ignoring ratings for −3 dB than for the clear (t_47_ = 6.125, p_Holm_ = 5.6·10^-9^, d = .892) and +6 dB conditions (t_47_ = 4.598, p_Holm_ = 6.7·10^-5^, d = .600), and for +6 dB than clear speech (t_47_ = 2.553, p_Holm_ = 0.014, d = .292; effect of Speech Clarity: F_2, 92_ = 24.187, p = 3.6·10^-9^, η_p_^2^ = .345). However, there was no effect of Distraction Type (F = .735, p = .482, η_p_^2^ = .016), nor an interaction (F_1, 46_ = 2.768, p = .103, η_p_^2^ = .057), indicating that participants were able similarly ignore speech under external and internal distraction.

Next, we calculated an rmANOVA – separately for the three TRF metrics – using the within-participants factors Speech Clarity (clear, +6 dB, −3 dB) and Attention (Attend, Ignore), and the between-participants factor Distraction Type (external, internal; Experiments 2 and 3). If there were any differences in the attention effect between distraction types, we would expect a significant interaction involving Attention and Distraction Type. In no analysis did we observe a significant interaction involving Attention and Distraction Type (for all p > 0.05), providing no evidence for attention effects to differ between the type of distraction.

## Discussion

The current EEG study investigated how disengagement from listening due to external and internal distractions alters neural speech tracking under different levels of background noise. We observed enhanced early neural responses (<0.2 s) to the speech envelope for speech masked by background babble compared to speech in quiet (Experiments 1-3), suggesting stochastic facilitation. Critically, neural speech tracking was reduced when individuals were distracted by a visual-stimulus stream (Experiment 2) or by internal imagination (Experiment 3). Overall, our study shows that disengagement from listening – towards external or internal spaces – that many people experience in crowded, noisy situations is associated with reduced neural encoding of speech.

### Background babble altered neural speech tracking

Neural speech tracking was assessed using different TRF metrics (Crosse et al., 2016): EEG prediction accuracy pools information across time (0-0.4 s), whereas P1-N1 and P2-N1 amplitudes indicate early sensory processing.

We observed a decline in EEG prediction accuracy with decreasing speech clarity (Figures 1-3), with the most prominent decline for the −3 dB SNR condition, mirroring story-comprehension accuracy and gist ratings (Figures 1-3). These results are consistent with previous research showing a close link between EEG prediction accuracy and speech perception (Ding and Simon, 2013; Vanthornhout et al., 2018; Synigal et al., 2023).

Early TRF responses showed a different picture. P1-N1 and P2-N2 amplitudes for speech at moderate and difficult levels of background noise were increased relative to clear speech. This increase may be surprising. However, several recent works observed enhanced early speech-tracking responses for speech presented in babble noise (Panela et al., 2024; Herrmann, 2025a, b). The findings are also in line with research using tone bursts, syllables, and high-frequency sound modulations, showing noise-enhanced early sensory responses (Alain et al., 2009; Parbery-Clark et al., 2011; Alain et al., 2012; Shukla and Bidelman, 2021).

Recent research specifically examined the noise-related enhancement of early neural tracking responses, showing that the enhancement is independent of attention, present at very high SNRs, and strongest for multi-talker babble (Herrmann, 2025a, b). These findings argue against a listening effort or attention mechanism, but instead suggest stochastic resonance/facilitation (Stocks, 2000; Stein et al., 2005; McDonnell and Ward, 2011). Background noise can introduce random fluctuations in membrane potential that transiently bring neurons closer to their firing threshold (Ward et al., 2002; Moss et al., 2004; Ward et al., 2010), effectively priming them to respond when speech-related input arrives. This stochastic resonance mechanism increases the probability of stimulus-driven, time-locked firing without inducing independent noise-driven firing (McDonnell and Abbott, 2009; McDonnell and Ward, 2011), thereby enhancing neural tracking of the speech envelope rather than degrading it through masking. Regardless of the specific mechanisms, the current study shows noise-related enhancements in neural speech tracking across all three experiments.

In addition to amplitude changes, the response morphology also appeared to vary with background noise, including a negative waveform shift (Figures 1B, 2E, 3C). Although morphology was not the focus here, the pattern may reflect a more fundamental change in the operating state of auditory system under noisy conditions. Prior work has shown that sustained noise can elicit ultra-low-frequency auditory-cortical activity (Ross et al., 2002; Okamoto et al., 2011; Barascud et al., 2016; Herrmann et al., 2022), though mechanisms remain incompletely understood. Background noise may therefore modulate both transient speech-envelope responses and the baseline state of auditory cortex.

### External and internal distraction reduces neural speech tracking

The main purpose of the current study was to examine how disengagement from speech listening due to external and internal distraction affects the neural encoding of speech as indicated by the TRF metrics. EEG prediction accuracy, P1-N1 amplitude, and P2-N1 amplitude all decreased when individuals attended to an external visual stream or internal thoughts and imagination.

Participants were able to disengage from listening in both the external and internal distraction tasks. Enhanced inter-trial phase coherence when attending to the visual stream, and ignoring speech, is consistent with previous work on attention and visual steady-state responses (Müller and Hillyard, 2000; Joon Kim et al., 2007; Keitel et al., 2017; Keitel et al., 2019), and with participants’ subjective reports of being able to ignore the stories well. Similarly during the imagination task, participants indicated being able to ignore and unable to retell the story. They also stated that they could visualize well mentally. Subjective reports are common measures of mental attentional drift (Smallwood and Schooler, 2006; Stawarczyk et al., 2011; Weinstein, 2018), because such internal changes resist objective assessments. The reported subjective experiences point to mental disengagement from listening during both distraction tasks.

Previous work on attentional disengagement from speech listening has mainly focused on speech in the presence of one competing speaker, finding that the tracking of the speech envelope of a target speaker is reduced when listeners disengage from listening to it (Ding and Simon, 2012; Power et al., 2012; Hambrook and Tata, 2019; Fuglsang et al., 2020; Kraus et al., 2021; Straetmans et al., 2024). Fewer works have shown attention effects when individuals attend to visual distractors while ignoring speech (Vanthornhout et al., 2019; for divided attention see Xie et al., 2023), or under ambient speech-masking conditions, such as multi-talker babble. Such conditions, however, reflect common scenarios in which listeners disengage in daily life, making them prime targets for understanding altered speech encoding. Attention effects in single competing-speaker paradigms have further been described as changes in TRF morphology, such as a reduced/absent P2 component for ignored speech (Ding and Simon, 2012; Fiedler et al., 2019; Kraus et al., 2021). By comparison, in the present babble-masking distraction paradigms, attention effects were expressed mainly as reductions in TRF metrics, with limited evidence for clear morphology-specific changes (Figures 2 and 3).

Importantly, the current work is first to show reduced speech encoding when individuals disengage from speech listening and engage in internal thought and imagination. This is critical, because internally oriented disengagement from listening is common among older people with hearing difficulties when listening effort exceeds available resources and/or perceived conversational benefit (Heffernan et al., 2016; Pichora-Fuller et al., 2016; Herrmann and Johnsrude, 2020). We used an intentional imagination task over a spontaneous mind wandering task as a proof on concept. The processes underlying intentional imagination and spontaneous mind wandering may be similar (Villena-González and Cosmelli, 2020; Safati et al., 2024), although others have argued that dissociating them may be beneficial (Forster and Lavie, 2009; Seli et al., 2016a; Seli et al., 2016b). Critically, the reduced speech tracking due to internal thought highlights neurophysiological feedback mechanisms, supressing auditory cortex activity in the light of mental imagination.

Our data provide little evidence that external and internal distractions differ in their suppression of neural speech tracking, suggesting that both may converge on a common pathway through which attention decouples from sensory processing. Although visual distraction engages visual sensory areas that internal thoughts may not, both likely recruit prefrontal and parietal networks that feed back to auditory cortex (Corbetta and Shulman, 2002; Braga et al., 2013; Puschmann et al., 2024), similarly reducing auditory-cortical gain and neural speech tracking. However, although no difference in the self-reported ability to ignore speech was observed, the external and internal distraction tasks were not explicitly equated for difficulty/cognitive load. Therefore, similar TRF reductions across experiments do not by themselves establish identical pathways, but could also reflect different mechanisms that produce a comparable degree of attentional withdrawal from speech.

Nevertheless, there were a few surface-level differences between external and internal distractions that may require additional research. Based on behavioral mind-wandering work (Steindorf et al., 2023) and consistent with behavioral ratings in the current study (Figures 2B and 3B), we had expected the effects of attention on neural speech tracking to be greatest under the challenging −3 dB SNR condition. This was observed for some TRF metrics during external visual distraction (e.g., Figure 2F), but not during internal distraction, for which we only observed a main effect of attention, but no interaction with speech clarity. This may potentially indicate that internal thought and imagination can lead to a blanket form of disengagement from the acoustic surroundings, whereas a distracting visual stream leaves more capacity for speech encoding in quiet, but very little when speech is presented in noise. Nevertheless, direct comparisons between distraction types did not show any significant differences. It is worth noting, however, that the current experimental design was not specifically suited for such direct interpretations due to its between-participant nature. This is thus an area for future research.

## Conclusions

This study investigated how disengagement from speech listening due to distractions by external visual stimuli or internal thought alters neural speech encoding under different degrees of background masking babble. We show that background babble enhances early neural responses to the speech envelope, consistent with recent reports of stochastic resonance. Critically, we show that external and internal distractions both lead to reduced neural tracking of the amplitude-onset envelope of speech, pointing to a converging neural pathway that supresses auditory cortex. The current study provides a neurophysiological marker that indexes different ways of disengagement from listening that are common among older adults with hearing challenges.

## Data availability

The data will be made available to others upon reasonable request.

## Acknowledgements

We thank Tiffany Lao and Saba Junaid for their support with stimuli creation and data collection. This study has been supported by BH’s Canada Research Chair program (CRC-2023-00383) and BH’s Natural Sciences and Engineering Research Council of Canada (Discovery Grant: RGPIN-2021-02602).

## Author contributions

**YR**: Methodology, data curation, formal analysis, visualization, writing - original draft, writing - review and editing, project administration. **MEC**: Data curation, formal analysis, writing - original draft, writing - review and editing, project administration. **BH**: Conceptualization, methodology, formal analysis, writing - original draft, writing - review and editing, supervision, funding acquisition.

