## Supplementary Materials for "Attentional disengagement during external and internal distractions reduces neural speech tracking in background noise"

### Analysis of Word Onset and Semantic Surprisal Responses

In addition to the analyses of the tracking of the envelope-onset provided in the main manuscript, we were interested in exploring whether semantic processing the speech is affected by external and internal distraction. One approach is to investigate how semantic surprisal is tracked by the brain over time (Brodbeck et al., 2018; Broderick et al., 2018; Broderick et al., 2021; Gillis et al., 2021; Broderick et al., 2022; Yasmin et al., 2023; Chalehchaleh et al., 2025).

We provide these analyses in the Supplemental Materials rather than in the main manuscript text, because it was unclear whether the responses can be confidently attributed to semantic processing per se.

#### Calculation of semantic word surprisal

For each word of each story, we calculated their semantic surprisal value using OpenAI's Davinci model (davinci-002). The Davinci model is the last model – with approximately GPT-3 capability – that enables the extraction of the word probability for each word, which, in turn, is the basis for calculating semantic surprisal for each word given previous context. Subsequent GPT models do not provide word probability any longer, although emerging powerful offline models like gemma-3-12b offer this. By using the Davinci model, the current approach capitalizes on a newer model than most previous approaches to extract semantic surprisal using GPT2, word2vec, GLoVe, or n-gram models (Brodbeck et al., 2018; Broderick et al., 2018; Broderick et al., 2021; Gillis et al., 2021; Broderick et al., 2022; Yasmin et al., 2023; Chalehchaleh et al., 2025). Table S1 gives an impression of the word surprisal values obtained by the Davinci model. We provide surprisal values for each word for two sentences, one with and one without a semantic violation at the final word (congruent vs incongruent), once with and once without further context:

| | | | | | | | | | | | | | | |
|---|---|---|---|---|---|---|---|---|---|---|---|---|---|---|
| ***Semantically congruent*** | | | | | | **The** | **landlord** | **drafted** | **the** | **motion** | **to** | **remove** | **the** | **tenant.** |
| | | | | | | - | 14.5 | 16.1 | 2.3 | 14.2 | 2.4 | 5.4 | 1.3 | 1.6 |
| **It** | **was** | **a** | **big** | **apartment** | **building.** | **The** | **landlord** | **drafted** | **the** | **motion** | **to** | **remove** | **the** | **tenant.** |
| - | 3.3 | 2.5 | 6.0 | 12.5 | 4.0 | 4.6 | 6.5 | 18.5 | 3.8 | 13.7 | 3.1 | 5.5 | 2.0 | 4.0 |
| ***Semantically incongruent*** | | | | | | **The** | **mechanic** | **replaced** | **the** | **part** | **to** | **fix** | **the** | **tenant.** |
| | | | | | | - | 16.6 | 10.5 | 0.6 | 6.5 | 8.0 | 2.4 | 0.5 | 19.1 |
| **The** | **car** | **shop** | **was** | **very** | **busy.** | **The** | **mechanic** | **replaced** | **the** | **part** | **to** | **fix** | **the** | **tenant.** |
| - | 10.6 | 14.0 | 4.4 | 16.1 | 2.7 | 3.2 | 5.5 | 8.6 | 0.8 | 5.1 | 8.5 | 2.5 | 0.7 | 20.3 |

**Table S1:** Semantic surprisal values for sentences that contain or do not contain a semantic violation (congruent vs incongruent) at the final word "tenant", once with and once without additional context.

The calculations provided in Table S1 show a large surprisal value for the semantic violation/incongruency at the final word “tenant”. Moreover, without context, the first few words of a sentence also exhibit high surprisal values, but this is reduced with context, showing surprisal is context sensitive. Hence, the Davinci model captures meaningful variation in semantic surprisal.

The onset time for each word in each story was obtained using Clarin’s forced alignment software (Schiel, 1999). Onset times were manually verified.

We used the word onset times to obtain two regressors for the TRF analyses. A word-onset regressor was calculated as an impulse vector that was 0 at non-word onset samples and the median surprisal value at samples corresponding to word onset times. We used the word onset regressor as a control condition to capture acoustic and any other processing associated with the word. For the semantic surprisal regressor, an impulse vector was created that was 0 at non-word onset samples, whereas at the word onset samples, the word-specific surprisal values were placed. Analyses focused on data from Experiments 2 and 3 because these experiments included the attention factor.

**Results of TRF analyses**

TRF analysis procedures mirrored those described in the main manuscript. We first calculated several multi-feature models to examine whether models comprising semantic surprisal increased EEG prediction accuracy beyond models including word onset. For both Experiment 2 and 3, there were no significant differences between three models: 1) Amplitude onset-envelope + word onset; 2) Amplitude onset-envelope + semantic surprisal; and 3) AWS - Amplitude onset-envelope + word onset + semantic surprisal (for all $p > 0.2$). Hence, adding semantic surprisal did not lead to a higher EEG prediction accuracy. These analyses provide little evidence that the semantic surprisal explains variance beyond the word onset.

Next, we calculated the TRF time courses separately for the word-onset and semantic-surprisal predictors. We focused on parietal electrodes and the 0.35-0.55 s time window, because previous research has shown semantic effects for these electrodes and this late time window (Hahne and Friederici, 2002; Friederici et al., 2004; Broderick et al., 2021; Broderick et al., 2022). Mean TRF time courses for Experiment 2 and 3 are shown in Figure S1 and S2, respectively. Time courses show a negative deflection around 0.4 s and topographies reveal a parietal distribution. This may suggest some word-level semantic processing.

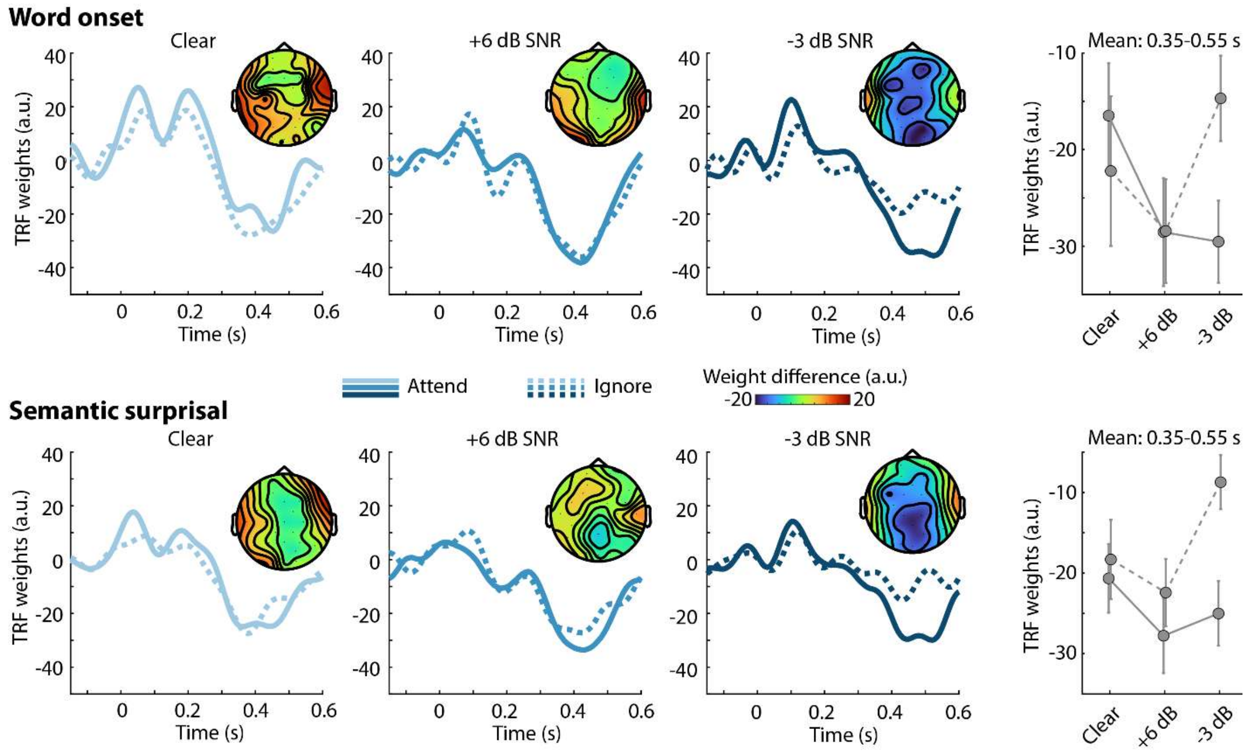


**Figure S1: TRF time courses and mean TRF weights for word-onset and semantic-surprisal predictors for Experiment 2. Top row:** TRF time courses (averaged across posterior electrodes) for speech-clarity and attention conditions for the word-onset predictor. Topographical distributions reflect the mean difference between ‘attend’ and ‘ignore’ conditions in the 0.35 s to 0.55 s time window. The plot on the right shows the mean TRF weights across the 0.35 s to 0.55 s time window. **Bottom row:** Same as top row for the semantic surprisal predictor.

A rmANOVA for the mean TRF weights in the 0.35-0.55 s time window was calculated separately for the word-onset predictor and the semantic-surprisal predictor, and separately for Experiments 2 and 3. Speech Clarity (clear, +6 dB, and -3 dB) and Attention Condition (attend, ignore) were used as within-participant factors.

For Experiment 2, the effect of Attention Condition was significant for the semantic surprisal predictor ($F_{2, 46} = 5.700$, $p = .026$), showing a more negative deflection for the attend compared to the ignore condition. None of the other effects or interactions were significant ($p > 0.05$).

For Experiment 3, for both the word-onset rmANOVA and the semantic-surprisal rmANOVA, the Attention Condition was significant ($F_{2, 46} > 6.5$, $p < 0.02$), showing a more negative deflection for the attend compared to the ignore condition. The Speech Clarity effect was also significant for both rmANOVAs ($F_{2, 46} > 7.5$, $p < 0.005$), in both cases due to the more negative deflection for the +6 dB condition compared to the other two conditions (for all $p_{Holm} < 0.05$; Figure S2, right). The interactions were not significant ($p > 0.05$).

Assuming the negative deflection around 0.4 s reflects some form of semantic word-level processing (even if not strictly due to semantic surprisal), the analyses may suggest that this word level processing is reduced under external and internal distraction, most consistently for the -3 dB SNR condition.

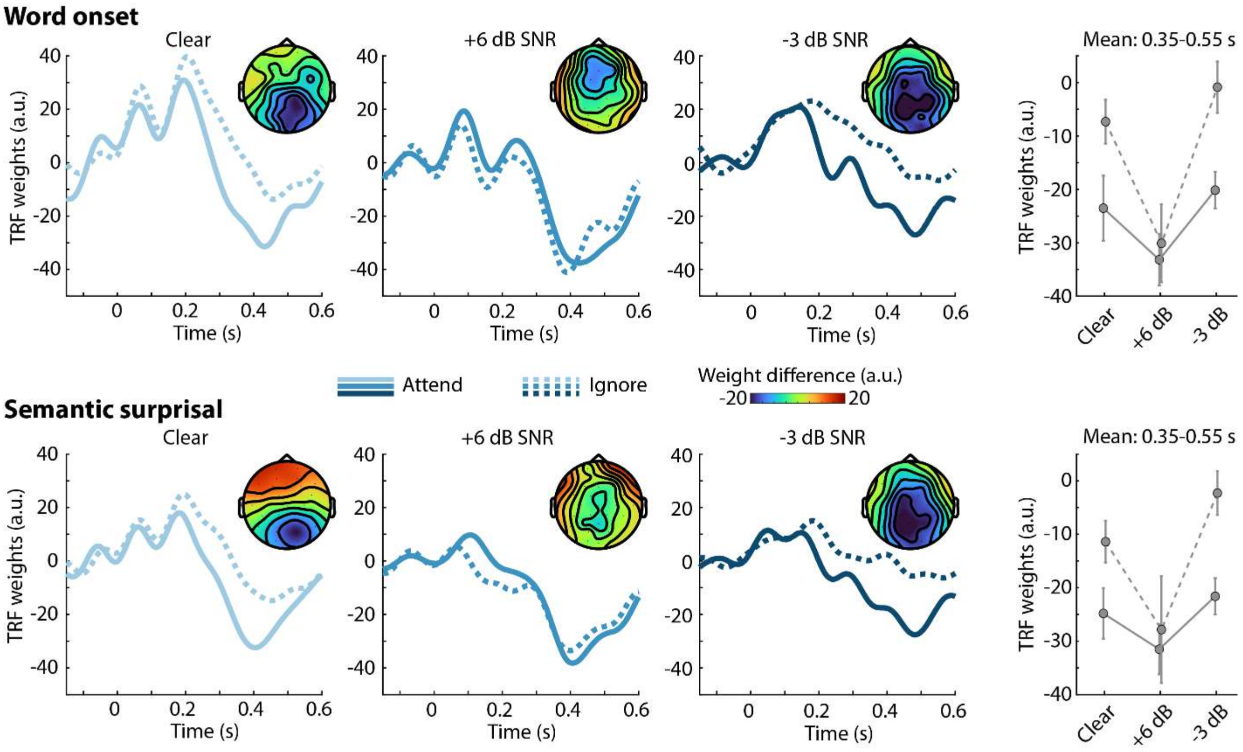


**Figure S2: TRF time courses and mean TRF weights for word-onset and semantic-surprisal predictors for Experiment 3. Top row:** TRF time courses (averaged across posterior electrodes) for speech-clarity and attention conditions for the word-onset predictor. Topographical distributions reflect the mean difference between 'attend' and 'ignore' conditions in the 0.35 s to 0.55 s time window. The plot on the right shows the mean TRF weights across the 0.35 s to 0.55 s time window. **Bottom row:** Same as top row for the semantic surprisal predictor.
